# Medial frontal cortex activity predicts information sampling in economic choice

**DOI:** 10.1101/2020.11.24.395814

**Authors:** Paula Kaanders, Hamed Nili, Jill X. O’Reilly, Laurence T. Hunt

## Abstract

Decision-making not only requires agents to decide what to choose, but also how much information to sample before committing to a choice. Previously established frameworks for economic choice argue for a deliberative process of evidence accumulation across time. These tacitly acknowledge a role of information sampling, in that decisions are only made once sufficient evidence is acquired, yet few experiments have explicitly placed information sampling under the participant’s control. Here, we use functional MRI to investigate the neural basis of information sampling in economic choice, by allowing participants to actively sample information in a multi-step decision task. We show that medial frontal cortex (MFC) activity is predictive of further information sampling prior to choice. Choice difficulty (inverse value difference) was also encoded in MFC, but this effect was explained away by the inclusion of information sampling as a co-regressor in the general linear model. A distributed network of regions across prefrontal cortex encoded key features of the sampled information at the time it was presented. We propose that MFC is an important controller of the extent to which information is gathered before committing to an economic choice. This role may explain why MFC activity has been associated with evidence accumulation in previous studies, in which information sampling was an implicit rather than explicit feature of the decision.

---

Decisions great and small – from food choices in a supermarket (Gidlöf et al., 2013) to selecting the next President (Nadeau et al., 2008) – are determined by the information that the decision-maker samples before they commit to a choice. Since the 1970s, cognitive psychologists have developed ways to determine how participants decide to sample information as a decision unfolds (Newell & Simon, 1972; Payne, 1976), which has led to a rich understanding of the role of information sampling in economic choice (Bettman et al., 1998; Hunt et al., 2016; Kobayashi et al., 2019; Krajbich et al., 2010; Navarro et al., 2016; Stewart et al., 2016). In such decisions, stimulus evaluation can influence subsequent information sampling, and vice versa. For example, attention to choice alternatives amplifies the value of the attended alternative (Smith & Krajbich, 2019), but a subjects’ currently preferred option also guides which information they sample next (Hunt et al., 2016; Shimojo et al., 2003). But perhaps the most pervasive effect of the value of choice alternatives on information sampling is that more difficult decisions take longer, providing more time for the agent to acquire information about the competing alternatives (Busemeyer & Townsend, 1993; Hunt et al., 2012; Jamieson & Petrusic, 1977; Milosavljevic et al., 2010).

Despite our rich understanding of the role of information sampling in economic choice, most studies of its neural basis have focused around simultaneously or subsequently presented choice items, without placing information sampling under the participant’s control (Gottlieb & Oudeyer, 2018). As a consequence, although neural mechanisms supporting information sampling are increasingly understood (Bisley & Goldberg, 2010; Blanchard et al., 2015; Horan et al., 2019; Stoll et al., 2016; Thompson & Bichot, 2005; White et al., 2019), much less is known about how future economic choices are guided by it. Nevertheless, accumulator frameworks often used to model economic choice in tasks where participants decide when to make a choice and terminate the trial themselves, tacitly acknowledge that decisions with longer reaction times are those in which more information is sampled. For example, perceptual and economic choice are often described using a drift diffusion model (DDM) of two-alternative forced choice, in which evidence is accumulated over time and integrated until one of two response boundaries is reached (Krajbich et al., 2010; Ratcliff & McKoon, 2008; Usher & McClelland, 2001). Implicit in the DDM, and other accumulator models of decision-making, are sequential choices of the agent to sample more information. If the bound is not yet reached, the agent continues to sample further information either from the environment or from memory (Shadlen & Shohamy, 2016).

Neural implementations of the DDM or other accumulator models have been used to identify brain networks that show similar trial-by-trial changes in aggregate activity as the model, and therefore could be evidence accumulators guiding choice. However, it is also possible that the trial-by-trial aggregate accumulator fluctuations may reflect greater information sampling on these trials. A region of medial frontal cortex (MFC) that encompasses the dorsal anterior cingulate cortex (dACC) and the adjacent pre-supplementary motor area (preSMA) has been identified as correlating with aggregate accumulator activity in a number of human decision-making studies using blood oxygen level dependent (BOLD) fMRI (Gluth et al., 2012; Hare et al., 2011; Pisauro et al., 2017; Rodriguez et al., 2015; Venkatraman et al., 2009; Zhang et al., 2012). Typically, aggregate accumulator activity is greatest on trials where choice difficulty (i.e. inverse value difference between alternatives) is highest. Yet because none of these studies explicitly measured information sampling on a trial-by-trial basis, it is unclear which of these two accounts – choice difficulty or information sampling – better explains BOLD responses in these areas (although importantly they are not mutually exclusive). Choice difficulty itself has also previously been associated with activity in MFC BOLD fMRI signal (Grinband et al., 2006; Pochon et al., 2008; Shenhav et al., 2014), as have reaction times in decision tasks (Thielscher & Pessoa, 2007). Yet on the other hand, this area has also been studied in the context of the exploitation-exploration dilemma, where MFC activity is linked to decisions to sample more information or explore new alternatives (Badre et al., 2012; Blanchard & Gershman, 2018; Boorman et al., 2009; Daw et al., 2006; Domenech et al., 2020; B. Y. Hayden et al., 2009; Kolling et al., 2012).

Here, we used BOLD fMRI to investigate the neural basis of information sampling in economic choice, by allowing participants to actively sample information in a multi-step decision task. Our experimental paradigm mirrors equivalent recent paradigms used in non-human primates (Hunt et al., 2018) and a large-scale human behavioural study (Hunt et al., 2016). As expected, subjects sampled more information when decisions were more difficult. A broad network of brain regions signalled the value of the presented cues, as indexed using representational similarity analysis (Hunt et al., 2018). When participants were first able to make a choice, fMRI signal within MFC correlated with the difficulty of the decision. Crucially, this effect was significantly reduced by including a co-regressor indicating whether the subject sampled further information prior to choice. Activity in MFC (and bilateral intraparietal sulcus) was instead explained by the decision to sample further information.

## Results

30 healthy human participants (aged 18-50) performed an information gathering and economic choice task inside the MRI scanner (Figure 1A). In the task, they were asked to choose between two pairs of cues, where the pair of cues on the left were one choice option and the pair on the right another. Each pair consisted of a probability and a magnitude cue. The reward associated with an option was the number of points represented by the magnitude cue awarded probabilistically in accordance with the probability cue. The cues were pictures of faces and houses and the magnitude or probability associated with them was learned by the participants in a behavioural session that took place before the main experiment (Figure 1B). ‘Optimal’ behaviour in the task (maximizing long-run expected reward) would be to choose the option with the higher expected value (=reward probability * magnitude).

**Figure 1.**
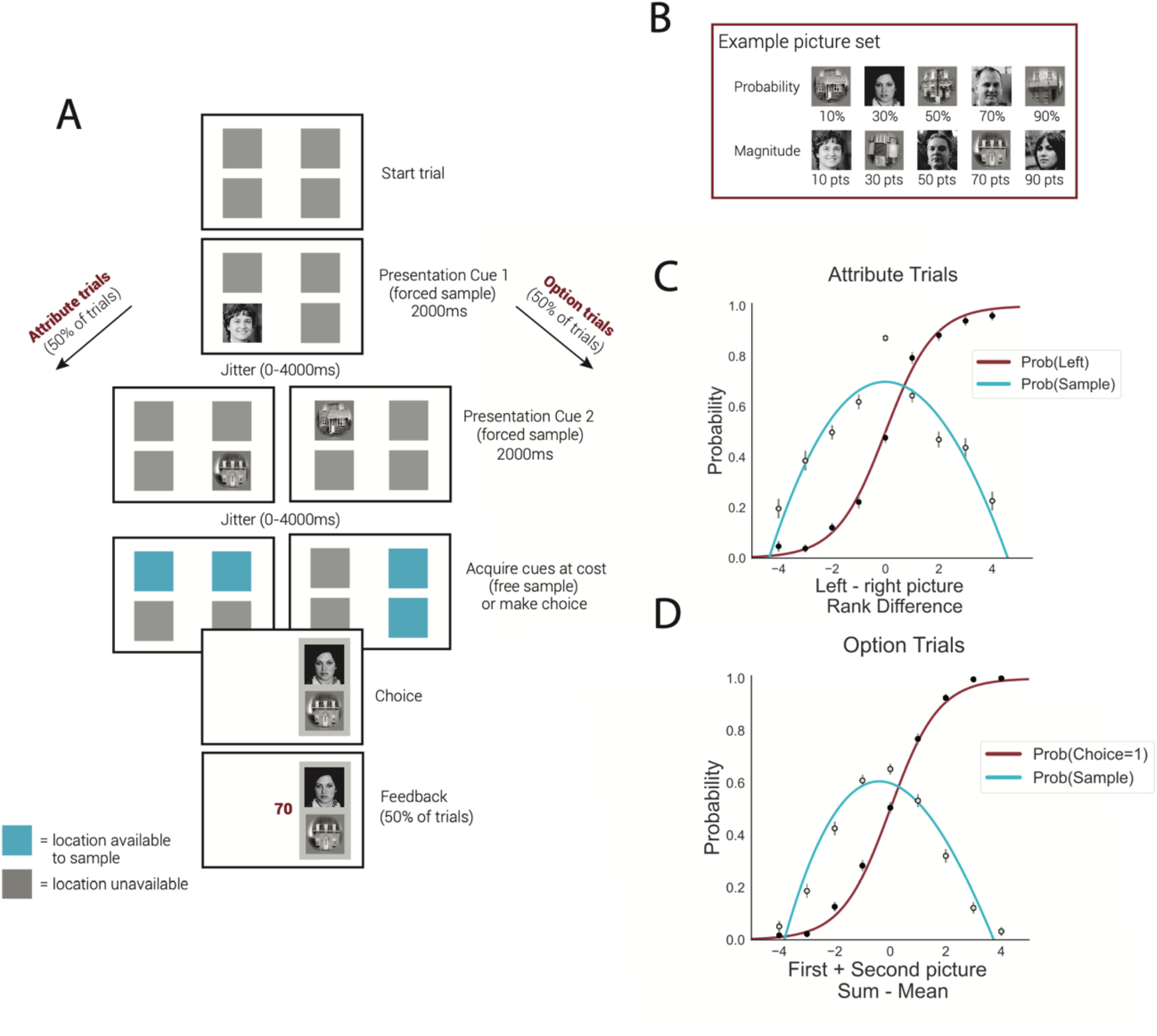
Experimental paradigm and participant behaviour. (A) Task structure. The task starts with all four cues being hidden. Two cues are shown sequentially. After this, participants can choose to sample more cues or make a choice between the two options. (B) Example picture set and their associated magnitude and probability amounts. (A,B) Note that the pictures of faces used in these panels do not belong to individuals and were generated. (C, D) Participants effectively use the cue values to make correct choices and to guide whether more cues need to be sampled. (C) In attribute trials, participants are more likely to choose the left option, the larger the value difference is between the left and right observed cues. Participants are more likely to sample additional cues the closer this value difference is to 0. These psychometric curves are plotting the probability of choosing the left option and sampling a third cue as a function of the difference in value rank between the two revealed cues. (D) In option trials, participants are more likely to choose the option of which the cues have been revealed, the higher the sum of these presented cue values is compared to the mean value of an option. Participants are more likely to sample additional cues the closer this value sum is to the mean. These psychometric curves are plotting the probability of choosing the option the cues of which were revealed and sampling a third cue as a function of the sum of the ranked values of the revealed cues subtracting the mean cue rank of an option.

Crucially, each trial started with the four cues being hidden. Participants were initially shown two of the hidden cues sequentially in a pseudorandom order determined by the experimenter. These could either be two cues from the same option (‘option trials’, 50% of trials) or two cues from different options representing the same attribute (probability or magnitude; ‘attribute trials’, 50% of trials). After these first two cues had been viewed, participants could choose whether to sample the remaining hidden cues by pressing the corresponding buttons and paying a small number of points (−3 points to sample the third cue, −6 points to sample the fourth cue). Alternatively, participants could forgo the opportunity to sample further information and instead make a choice between the two options based on the information sampled thus far. Subjects then received feedback on how many points they earned that trial, although in order to decouple this haemodynamic response from that of the decision onset, feedback was only delivered on half of all trials (Guitart-Masip et al., 2011).

Both participants’ eventual choices, and their propensity to sample more information prior to choice, depended upon the values associated with the first two presented cues. In attribute trials, their choices were a function of the difference in value between the left and right cues (t(29)=10.79, p<0.001; Figure 1C), whereas in option trials their choices depended upon the value of the option relative to the mean expected value (t(29)=15.88, p<0.001; Figure 1D). Importantly, participants’ decision to sample a third piece of information also depended upon these cue values, such that the more difficult the decision, the more information was sampled (t(29)=-2.11, p=0.04 on attribute trials; t(29)=-7.20, p<0.001 on option trials; Figure 1C-D). In attribute trials, the direction of this sample tended to be biased towards the option that currently had the highest value (Figures S1-2), a bias also observed in our previous monkey (Hunt et al., 2018) and large-scale human behavioural studies (Hunt et al., 2016).

We first performed a whole-brain searchlight representational similarity analysis (RSA), using templates representing several features of the task at *the time of presentation of the first cue* (Figure 2). At each 100-voxel searchlight sphere we regressed the templates onto the RSA matrix of that location to find the regions where brain activity showed a similar representation. Several of these templates were the same as those used in our previous study using single unit recording in macaques (Hunt et al., 2018). First, as a positive control for the success of our searchlight RSA pipeline, we included a face/house template which represented which category a cue belonged to (Figure 2A). Replicating numerous RSA results in the prior literature (Dima et al., 2018; Kriegeskorte et al., 2008), this produced a strong correlation with visual and inferior temporal cortex (peak MNI coordinates at [31, −61, −9] and peak Z=12.37, p<0.001, Figure 2C). Notably, these results were robust and replicable even at the level of individual participants (Figure S3). The next template identified regions the multivariate activity of which distinguished between whether the currently presented cue was on the left or right hand side of the screen (Figure 2C). As expected, this was strongly represented in primary visual cortex (peak MNI coordinates at [−6, −85, 4] and peak Z=11.28, p<0.001, Figure 2D). Surprisingly, however, we did not find activation in frontal eye fields or dorsolateral prefrontal cortex, despite this pattern clearly being represented in dlPFC in our previous single-unit data (Hunt et al., 2018).

**Figure 2.**
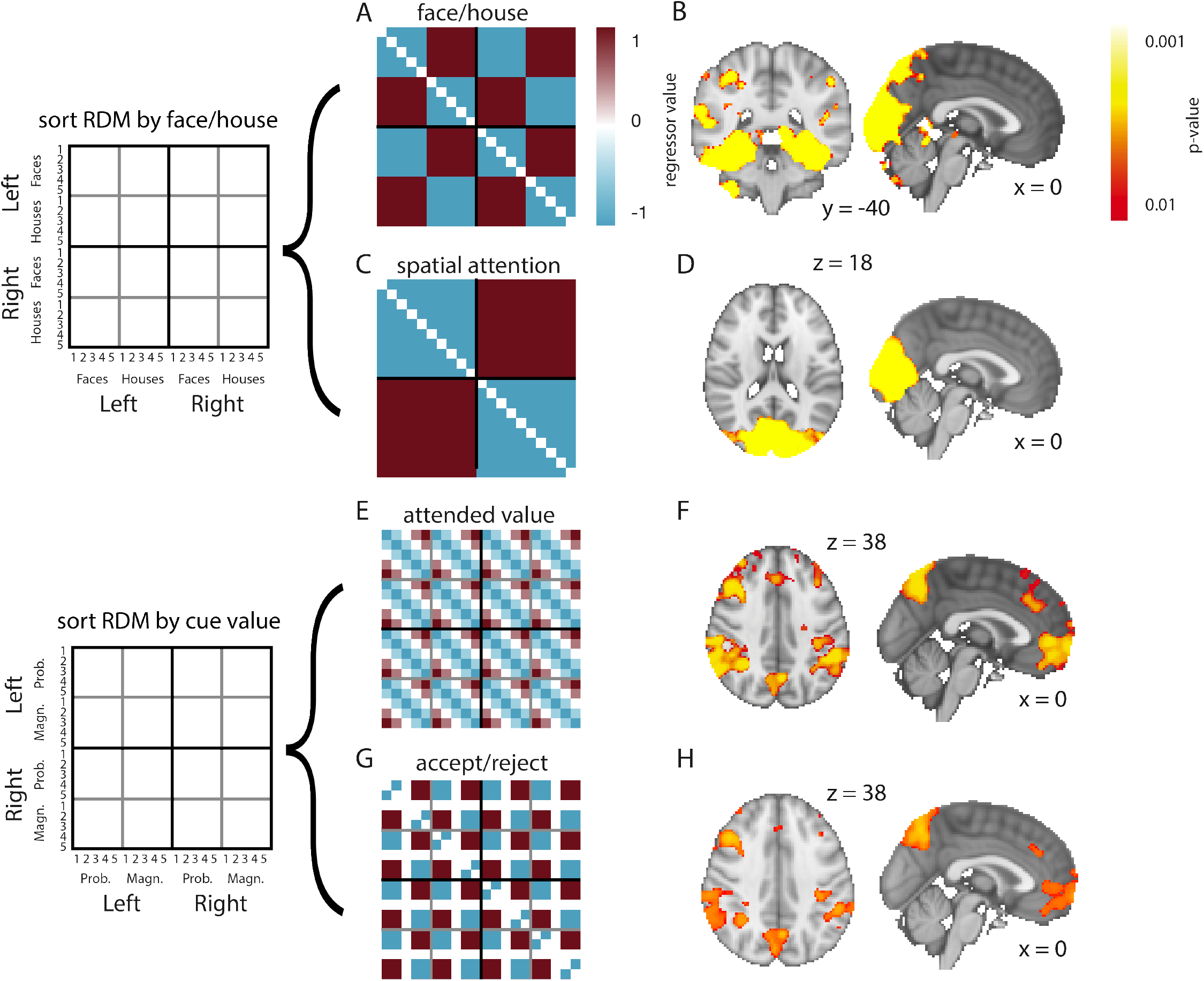
Whole-brain searchlight analysis reveals representations of several features of the task. (A,C,E,G) RSA templates for spatial attention, face/house category, attended value, and belief confirmation. (B) Cue category (face or house) is represented in visual and inferior temporal cortex. (D) Spatial attention is represented in visual cortex. (E-H) We find activations for attended value and belief confirmation in an elaborate network including vmPFC, ACC, and posterior parietal cortex. All parametric maps are depicting p-values derived from using threshold-free cluster enhancement (TFCE) on GLM z-score maps.

To investigate the neural correlates of the value-related features of the task, we included ‘attended value’ and ‘accept/reject’ templates, mirroring those in our previous study (Hunt et al., 2018). The ‘attended value’ template simply represents the value of the first cue, ranked from 1-5 and then normalized, such that low-value cues are predicted to be similar to other low-value cues, and high-value cues are predicted be similar to other high-value cues (Figure 2E). The template assumes that magnitude and probability cues were treated the same in terms of how valuable they were, because to calculate the expected value of an option participants had to collapse across the two cues, and because magnitude and probability cues were found to be similarly encoded in macaque OFC single-cell recordings (Hunt et al., 2018). The ‘accept/reject’ template also represents the value of the first cue but binarises this into ‘above average’ or ‘below average’ in value (or ‘neutral’, for the mid-value cue; Figure 2G). In our single-unit data, this correlated with activity in dACC (Hunt et al., 2018). In our fMRI data, we found that both the ‘attended value’ and ‘accept/reject’ templates correlated with the activity in an extensive network within frontal and parietal cortex, including ventromedial PFC (vmPFC; peak at MNI coordinates [2, 50, −1] and peak Z=4.36, p<0.001 for ‘attended value’ template; peak at MNI coordinates [2, 51, 1] and peak Z=3.90, p=0.006 for ‘accept/reject’ template), dACC (peak at MNI coordinates [0, 37, 29] and peak Z=4.80, p=0.003 for ‘attended value’ template; peak at MNI coordinates [0, 35, 31] and peak Z=5.91, p=0.006 for ‘accept/reject’ template) and posterior parietal cortex (peak at MNI coordinates [52,-49,51] and peak Z=5.91, p<0.001 for ‘attended value’ template; peak at MNI coordinates [52, −49, 51] and peak Z=5,14, p=0.002 for ‘accept/reject’ template; Figure 2E-H). As expected, this network corresponds well with areas previously identified in meta-analyses of subjective value encoding using fMRI (Bartra et al., 2013; Clithero & Rangel, 2014).

We then investigated the neural correlates of the decision to sample further information, which first occurred at the time of presentation of the second cue, and whether this might explain activations that would otherwise be described as encoding choice difficulty. To test this, we used a mass univariate analysis, as the task comprised too many conditions to perform RSA at the time of presentation of the second cue. In an initial analysis, we included only the values of the different cues as regressors, and found a subregion of MFC that was negatively related to the difference between the values of the chosen and unchosen options, which we will refer to as ‘inverse value difference’ (Figure 3A, peak at MNI coordinates [−2, 14, 52] and peak Z=5.37; see uploaded maps at neurovault.org for unthresholded Z-statistic maps for this and all other analyses). In other words, activity in this region was greater for more difficult decisions, replicating a large number of previous findings in the economic choice literature (Grinband et al., 2006; Pochon et al., 2008; Shenhav et al., 2014).

**Figure 3.**
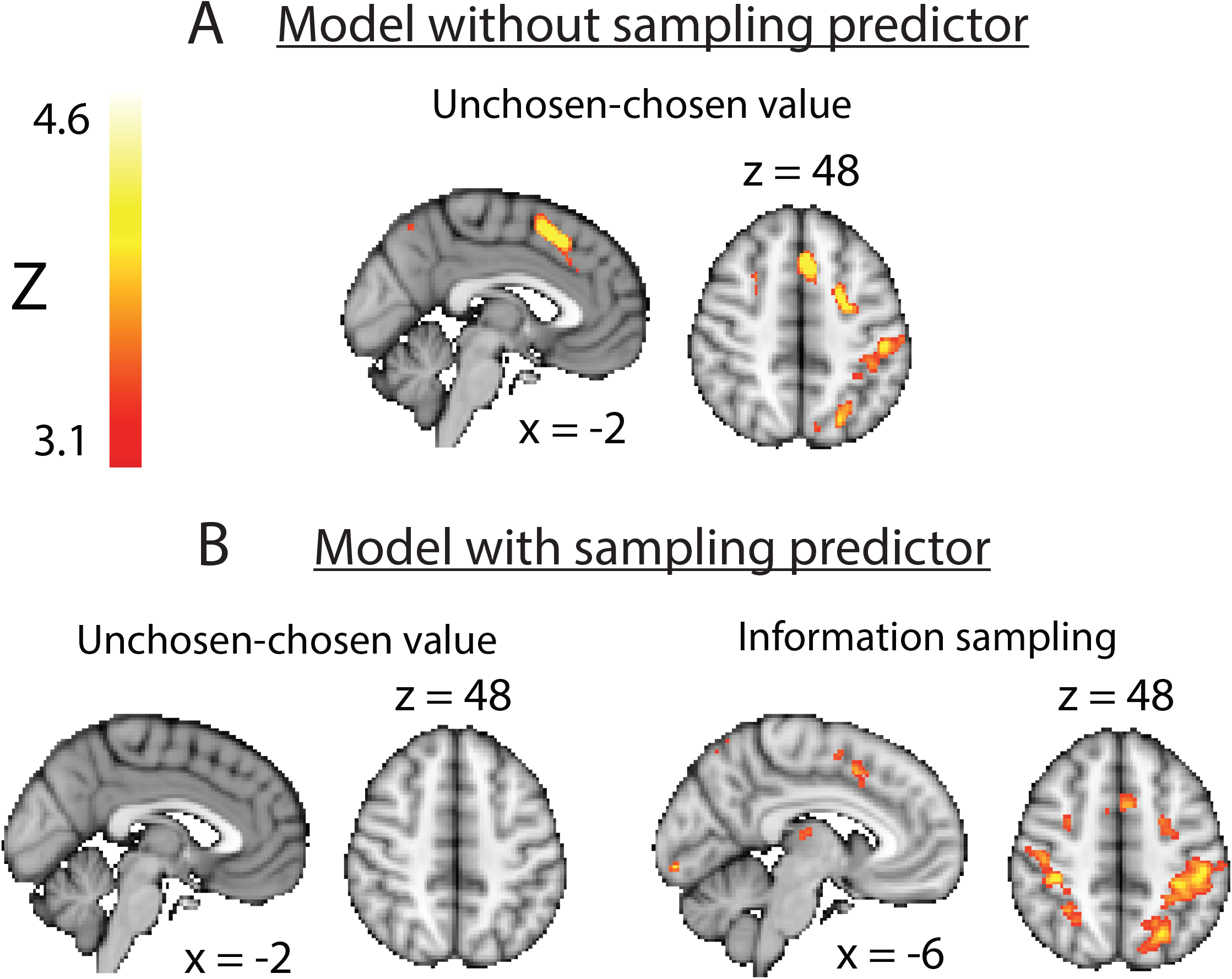
There is a main effect of inverse value difference (choice difficulty) at the time of presentation of cue 2 in MFC, which disappears when subsequent information sampling is included as a regressor. (A) Linear positive effect of inverse value difference in MFC on option trials in 30 subjects. (B) There is no effect of inverse value difference in MFC when information sampling is included as a regressor, while there is a linear positive effect of information sampling in MFC in 24 subjects. All parametric maps are cluster-corrected and thresholded at z>3.1.

Crucially, this model did not yet account for whether participants chose to sample further information before committing to one of the options. In a further regression model, we included a binary information sampling regressor describing whether or not participants chose to sample more information on a given trial. As expected from the relationship between difficulty and information sampling (Figure 1C/D), this regressor had some correlation with the regressor encoding value difference (average correlation of r=-0.3; Figure S5; we note that 5 participants were excluded from this analysis, as they sampled additional information before choosing less than once per block). We found that the main effect of choice difficulty in MFC disappeared when the information sampling predictor was included, and this was not a consequence of the exclusion of the 5 participants who rarely sampled further information (Figure S8). Instead a main effect of information sampling was seen in a small section of MFC, which was more active on trials where subjects decided to sample further information (Figure 3B, Figure 4, peak activation at MNI coordinates [−6, 10, 46] and peak Z=4.04). Consistent with recent reports that highlight the role of parietal cortex in reducing uncertainty about future rewards (Horan et al., 2019), we also found bilateral activation of intraparietal sulcus for this regressor (peak activation at MNI coordinates [−36, −32, 42] and peak Z=5.78, Figure 3B).

**Figure 4.**
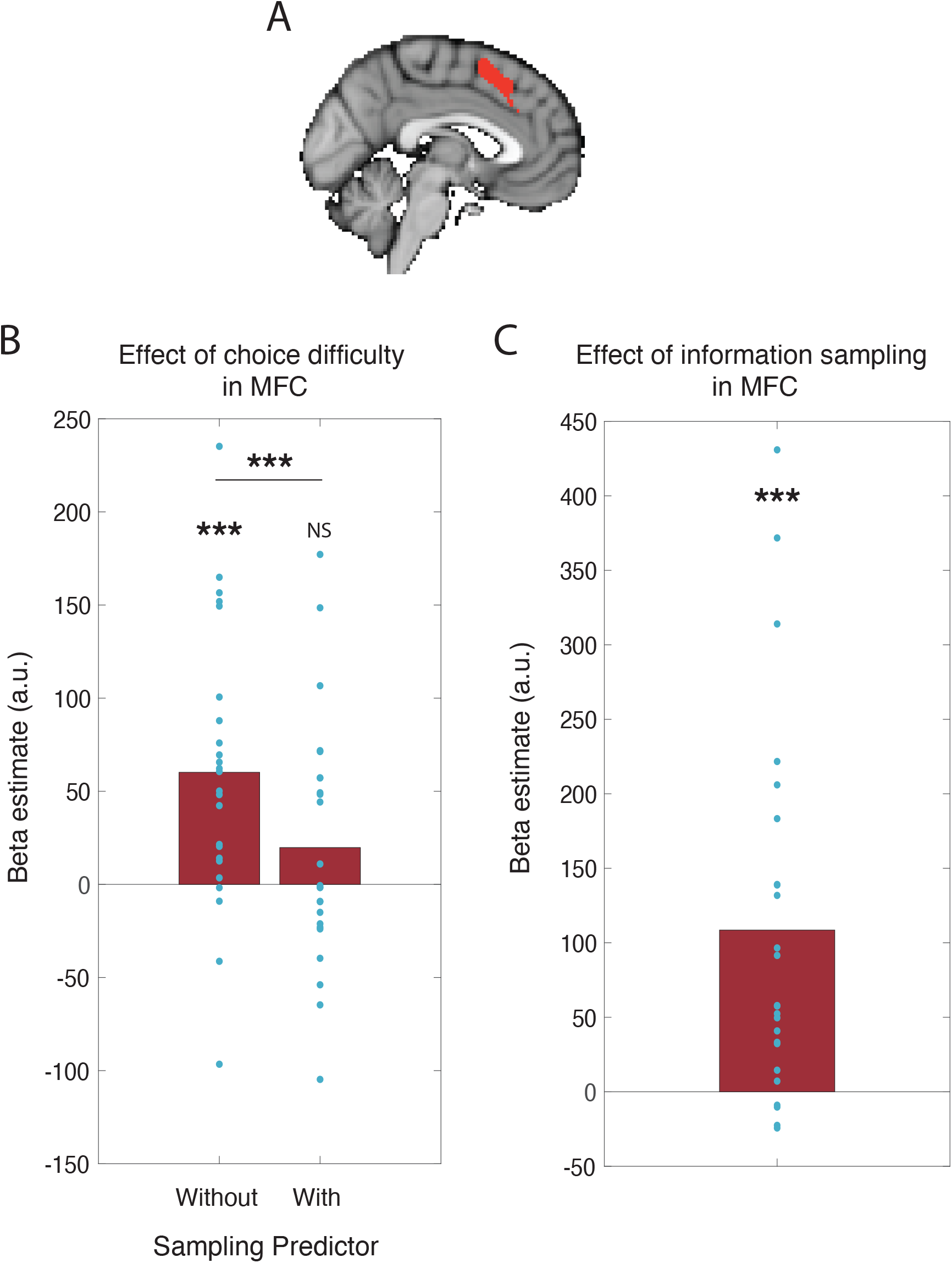
When information sampling is included as a co-regressor here is a significant decrease in the main effect of inverse value difference (in an MFC region of interest equivalent to the region defined in Figure 3A, defined by a leave-one-out procedure). (A) MFC region of interest derived from mass univariate analysis including all 30 subjects. (B) Effect of inverse value difference in MFC in GLMs with and without the information sampling co-regressor in 24 subjects. (C) Effect of information sampling in the same brain region. A stronger BOLD signal was found in this region on trials where participants would subsequently decide to sample more information before committing to a choice compared to trials where they made a choice straight away.

To further explore whether choice difficulty encoding was significantly reduced by the inclusion of the information sampling regressor, we performed a region-of-interest analysis. We focused our results on the MFC region that encoded inverse value difference (i.e. choice difficulty) in the whole brain analysis (Figure 3A), but in order to avoid circularity (Kriegeskorte et al., 2009), we used a leave-one-out procedure in which each participant’s ROI was defined by taking significant MFC voxels from a group model including all participants except them (Boorman et al., 2013). This revealed a significant reduction in the encoding of inverse value difference as a consequence of including the information sampling regressor (Figure 4B; t(23)=3.28, p=0.003), such that choice difficulty was no longer significant encoded in this region (t(23)=1.17, p=0.26). Instead, this region of interest significantly encoded the decision to sample further information (Figure 4C; t(23)=4.17, p<0.001).

Finally, we investigated encoding of belief confirmation, which was defined as the extent to which the value of the second cue confirmed the belief formed after presentation of the first cue. In our single unit study, we found that belief confirmation was reliably encoded in dACC (Hunt et al., 2018), and similarly in the present study we found fMRI activity in a more anterior portion of MFC encoded belief confirmation on ‘option’ trials (Figure S4; peak Z=4.81 at peak at MNI coordinates [8, 34, 8]). However, this effect did not survive whole-brain correction in the reduced sample of 24 participants who regularly sampled more information before choosing; we therefore did not investigate further whether this regressor was significantly reduced by the inclusion of information sampling as a co-regressor.

## Discussion

Here we have shown that in an information sampling and economic choice task, MFC activity encodes whether a subject will sample further information before committing to a decision. We moreover found that this effect could explain away the encoding of choice difficulty (inverse value difference) in MFC, a signal that has been commonly observed in many previous studies of economic choice (Grinband et al., 2006; Pochon et al., 2008; Shenhav et al., 2014). Our findings should not be interpreted as suggesting that difficulty is not encoded in MFC, but instead that difficulty be a fundamental determinant of whether further information is sampled prior to choice. This forms a logical extension of previous work on the role of MFC as an evidence accumulator (Gluth et al., 2012; Hare et al., 2011; Pisauro et al., 2017; Rodriguez et al., 2015; Venkatraman et al., 2009; Zhang et al., 2012), as evidence accumulation involves an implicit decision to sample further information when a decision bound has not yet been reached (Shadlen & Shohamy, 2016). An advantage of the current information gathering paradigm is that it allowed us to study the relationship between MFC signals and subsequent behavior; most paradigms in the field are much simpler and do not allow for this type of analysis.

Our findings are consistent with previous studies that propose a role for MFC in guiding exploratory behavior (Badre et al., 2012; Boorman et al., 2009; Daw et al., 2006; Domenech et al., 2020; B. Y. Hayden et al., 2009; Kolling et al., 2012). For example, one recent study by Blanchard and Gershman (2018) contrasted trials in which participants decided to place a bet, versus trials where they decided to obtain further information about the value of that bet. Activity in MFC and bilateral insula was significantly greater on observe trials than on bet trials (we also found subthreshold activity in insula predictive of information sampling, see Figure S9 and maps on neurovault.org). Similarly, a single unit recording study by White et al. (2019) identified dACC as the leading node in a network of areas that encoded gaze shifts to resolve uncertainty about upcoming rewards, and further studies have shown that activity in this area predicts ‘checking’ behaviours when monkeys are close to receiving a reward (Stoll et al., 2016). On the other hand, it has also been suggested that dACC is particularly active when participants switch away from a default behavior (E. D. Boorman et al., 2013; Kolling et al., 2016). It is possible that we find mediofrontal activity predictive of information sampling because in our study not sampling additional information could be considered a default behavior, given that participants commit to a choice straight away more often than they decide to sample more cues (Figure 1C/D).

More broadly, different subregions of MFC have been associated with diverse parameters in different paradigms (Kolling et al., 2016), whether it is sampling value (Stoll et al., 2016), conflict (Botvinick et al., 1999), choice difficulty (Grinband et al., 2006; Pochon et al., 2008; Shenhav et al., 2014), or belief (dis)confirmation (Boorman et al., 2013; Hunt et al., 2018; Quilodran et al., 2008; Wessel et al., 2012). It is notable that all of these task parameters may affect the degree to which additional information sampling is required before committing to a choice.

Our behavioural results also indicate that humans are more likely to sample information from an option that they believe is good (Figure S1-2), consistent with our previous behavioural findings (Hunt et al., 2016). We hypothesize that MFC may therefore drive information sampling through active hypothesis testing (Markant & Gureckis, 2012). In other words, rather than sampling information from all available alternatives equally, the agent has a hypothesis that one of the alternatives is best, continues to sample from it and only moves to another alternative when the first alternative turns out to be a bad option. This is consistent with the presence of a ‘belief confirmation’ signal in ACC, which we find here (Figure S4) and also found in our previous single unit study (Hunt et al., 2018). It is also consistent with recent behavioural findings in perceptual decision-making (Talluri et al., 2018), and the recent suggestion that gaze affects the integration of goal-relevant evidence rather than value in economic choice (Sepulveda et al., 2020). Such an account of economic choice has recently been proposed as particularly ecologically valid in decisions containing multiple options (Hayden, 2018). More detailed testing of this hypothesis will require paradigms in which multiple options are available for choice, and where information sampling and choice are explicitly dissociated from one another.

We were able to identify brain networks encoding both sensory- and value-related aspects of the task using RSA, some of which encompassed areas in PFC (Figure 2) whose value responses have previously been shown using multivariate pattern analysis (Kahnt et al., 2010). A similar region was found to encode continuous and categorial (‘below’ or ‘above’ average) attended value. We tested for the presence of these two regressors despite their correlation because in the macaque single-cell data from Hunt et al. (2018), continuous value was found to be mostly encoded in OFC, while categorical value was encoded in ACC. It is possible that we were not sensitive to detect a similar dissociation here due to the lower signal-to-noise in human fMRI data compared to that in single-cell data. In the same analysis, we also observed value-related activity in bilateral parietal cortex, which is consistent with a similar RSA regressor used to describe parietal cortex EEG responses to numerical magnitude in an economic choice task (Luyckx et al., 2019).

The clear correlation between templates describing the visual properties of cues in the task and visual/inferotemporal cortical activity replicates earlier work showing that RSA distinguishes objects of different categories in these areas in human fMRI data (Dima et al., 2018; Kriegeskorte et al., 2008; O’Toole et al., 2005). It has previously been suggested that such an approach can be used to study representations across species, found using different types of data (Kriegeskorte et al., 2008). We hypothesized that our task might allow us to perform a similar cross-species analysis in order to identify functional homologues of the PFC subregions studied with single unit recordings in Hunt et al. (2018). We did not clearly identify the same triple dissociation that was seen in our previous study (Figure S6, S7). It still remains unclear whether RSA really probes fine-grained spatial information that is similar to that identified by single neuron data, or whether it is primarily driven by the same macroscopic signals that can also be isolated in mass univariate analyses (Op de Beeck, 2010). It is possible that the intermixed positive and negative coding that supported the successful RSA in Hunt et al. (2018) is not observable at the voxel level in human fMRI. This may also explain why decoding accuracies in multivariate pattern analyses are invariably far lower in prefrontal cortex than in occipital and temporal cortices (Bhandari et al., 2018).

In summary, we have shown that MFC activity in economic choice predicts subsequent information sampling. We replicated previous studies showing that MFC activity in different subregions is related to choice difficulty, but critically this effect was explained away by the inclusion of information sampling as a co-regressor. This suggests that the role of MFC may extend beyond that of evidence accumulation, and implies an important role for MFC in guiding adaptive information sampling during economic choice.

## Methods

30 healthy human participants (aged 18-50) attended two study sessions: one behavioral training session (1hr) and one fMRI session (2hr 15min). One participant was excluded from the representational similarity analyses as their fMRI study session was ended prematurely, meaning we did not obtain enough data from this participant to appropriately balance across different conditions. In the behavioral session, participants learned the meanings of ten stimuli: five faces and five houses. Five of these stimuli represented a reward probability (10%, 30%, 50%, 70%, or 90%) and the other five a reward magnitude (10, 30, 50, 70, or 90 points). We ensured that whether the cues were faces or houses was orthogonal to cue values by pseudo-randomizing the meaning of each cue for each participant. Specifically, there were always 3 face probability cues and 2 house probability cues (and similarly for magnitude), or vice versa, and these were interspersed such that faces or houses could not be associated with generally low or high value (Figure 1B). After learning the cues, participants were trained on the main task.

In the main task, participants were asked to choose between two pairs of cues (Fig. 1A). Each pair consisted of a probability and a magnitude cue. The reward associated with that option was the number of points represented by the magnitude cue awarded probabilistically in accordance with the probability cue. ‘Optimal’ behavior in the task (maximizing long-run expected reward) would be to choose the side with the higher expected value (reward probability multiplied by reward magnitude). However, a trial started with the four cues being hidden (presented as grey squares), where the pair of cues on the left was one option and the pair on the right another. Participants were initially shown two different cues sequentially, the selection of which was pre-determined by the experimenter in order to balance cue presentation for representational similarity analyses. These could either be two cues from the same option (‘option trials’, 50% of trials) or two cues from different options representing the same attribute (probability or magnitude; ‘attribute trials’, 50% of trials). Note that the location of each attribute was counterbalanced to be the top or bottom cue, but that this was consistent between the two options within a trial (eg. if the top cue of the left option was the probability cue, the top cue of the right option was also the probability cue). The location and identity of the first cue were counterbalanced throughout each block. The reason the cues were shown sequentially is because this enabled us to study brain activity in response to each cue separately. At the second cue, we can also study choice difficulty and belief confirmation by looking at how the neural response depends on the value difference between the options and on how well the evidence provided by the second cue conforms to that provided by the first cue. Each cue presentation lasted 2s and was followed by a 0-4s jittered interval during which all the cues were hidden again.

After this, participants were given the opportunity to view the remaining two hidden cues by pressing the corresponding buttons. However, participants had to pay points to do this: 3 points to see a third cue, and another 6 points to see the fourth and final cue. As such, participants had to judge how useful the additional information provided by the last two hidden cues might be in ensuring they choose the option with highest expected value. In a pilot study, it was found that these costs typically led participants to sample further information on a subset of trials (cf. Fig. 1C/D; although we note that the propensity to sample further information still varied across individuals). Note that participants could not choose to see the first two cues again. When participants finished sampling the additional cues, or decided they did not want to sample further cues at all, they made a choice between the two options. On half of all trials, subjects received feedback on how many points they earned that trial; feedback was only revealed on 50% of trials, as this made sure stimulus-locked value-related signals and feedback-locked reward-related signals were decorrelated in the design matrix. Inter-trial interval was jittered and between 3-7s. Participants were only invited to take part in the fMRI part of the experiment if they chose the highest valued option on 70% of trials in the last practice block of the main task.

In the fMRI session, participants first received further training on the task by doing two blocks of 40 trials each outside the scanner. Inside the scanner, participants did 10 practice trials and five blocks of the main task (a total of 200 trials). Participants received £1 for every 300 points earned in the task, in addition to a £20 show-up fee for coming to both study sessions. A counter in the right bottom corner of the screen kept track of the money earned up to that point throughout the task. As such, participants earned an average of £41 overall. This experiment was realised using Cogent 2000 developed by the Cogent 2000 team at the FIL and the ICN and Cogent Graphics developed by John Romaya at the LON at the Wellcome Department of Imaging Neuroscience.

### fMRI Data Collection

Whole-brain fMRI measurements were made using a Siemens Prisma 3T scanner with a 2 x 2 x 2mm voxel size, repetition time (TR) = 1.235s, echo time (TE) = 20ms, flip angle = 65° with an axial orientation angled to AC-PC using a 64-channel head coil. The sequence used was MB3 PAT2. Participants performed five blocks of the main task inside the scanner with short breaks in between. For each new block a new run was initiated. Given that the task was not speeded, the runs were of variable length and so were the number of volumes collected per run. The average number of volumes collected per participant was 2587. T1-weighted structural images were obtained using an MPRAGE sequence with 1 x 1 x 1mm voxel size, on a 174×192×192 grid, TE = 3.97ms, TR = 1.9s. A field map with dual echo-time images was also acquired (TE=7.38ms, whole-brain coverage, 2.4 x 2.4 x 2.4 mm voxel size).

### fMRI Data Analysis

fMRI analysis was carried out using the FMRIB Software Library (FSL), and custom-written RSA scripts in MATLAB that built upon the Representational Similarity Analysis toolbox (Nili et al., 2014).

The pre-processing performed on the data was the following: motion correction using MCFLIRT (Jenkinson et al., 2002); non-brain removal using BET (Smith, 2002); spatial smoothing using a Gaussian kernel of FWHM 5mm; grand-mean intensity normalisation of the entire 4D dataset by a single multiplicative factor; highpass temporal filtering (Gaussian-weighted least-squares straight line fitting, with sigma=50.0s). Registration to high resolution structural and/or standard space images was carried out using FLIRT (Jenkinson et al., 2002; Jenkinson & Smith, 2001).

### Behavioral data analysis

We studied the effects of cue value on choice and sampling decisions using single-subject logistic regressions followed by one-sample t-tests on the single-subject regression coefficients (Figures 1C/D). Specifically, we performed two logistic regressions on attribute trials where the independent variable was the value rank difference between the left and right presented cues and the dependent variable was whether or not the left option was chosen or whether a third cue was sampled, respectively. Similarly, on option trials two logistic regressions were executed where the independent variable was the summed cue ranks of the two presented cues minus the mean value rank of an option and the dependent variable was whether or not the fully revealed option was chosen or whether a third cue was sampled, respectively.

### Representational Similarity Analysis (RSA)

We first estimated a whole-brain general linear model (GLM) with regressors encoding onsets of the first two cues, cue category (face or house) and what side of the screen the cue appeared on. This GLM was executed on the spatially smoothed data. Then, we performed searchlight RSA, meaning RSA is performed for a 100 voxel sphere of voxels, centered around each voxel in the brain (Nili et al., 2014). These were then regressed against model representational similarity matrices (Figure 2) to find areas of the brain the activity of which showed similar representations. We then performed a statistical test at each voxel and significant clusters were identified using threshold-free cluster enhancement (TFCE; Smith & Nichols, 2009). The resulting p-values were family-wise error rate controlled at a threshold of p<0.05. This same method was used with representational similarity matrices obtained from Hunt et al. (2018) to investigate similarities between representations in monkey and human prefrontal cortex, although here we present unthresholded statistical maps as no clusters survived multiple comparisons correction in PFC.

### Mass Univariate Analysis

Two GLMs were fit to the individual subject data, one including and one excluding regressors describing information sampling behavior. We also included regressors describing onset of the first two cues, cue values (separated into cues where the associated option was eventually chosen or unchosen), side of cue presentation, trial type (option or attribute trial), whether feedback was presented, and if so, how much reward the participant received on the preceding trial. Lastly, our model included a regressor that reflected belief confirmation, encoded as an index defining how much the second cue should change the agent’s belief about what the best option is (Figure S4). We used contrasts of parameter estimates to find regions that encoded the inverse value difference between the revealed cues (i.e. unchosen minus chosen value, or choice difficulty). All regressors were modelled as 2s boxcar functions, to match the duration for which the cue was presented. We report mass univariate results at the group-level using whole-brain family wise error (FWE) corrected statistical significance and a cluster significance threshold of Z>2.3 and p<0.05. Note that similar results were obtained using TFCE (Smith & Nichols, 2009).

To measure how much the main effect of choice difficulty was reduced by adding the information sampling regressor (Fig. 4), without being biased by the initial GLM, we used a leave-one-out procedure. BOLD activity was extracted for each participant from the choice difficulty (in both GLMs) and information sampling contrasts using an ROI defined by a group model excluding that participant. The ROI was defined as any positively significant clusters (Z>3.1, p<0.05, corrected) for the contrast of inverse value difference found within a large mediofrontal mask. We then performed a one-sample t-test on the extracted parameter estimates for the inverse value difference contrast, both including or excluding information sampling behaviour as a co-regressor.

---

**Figure S1.**
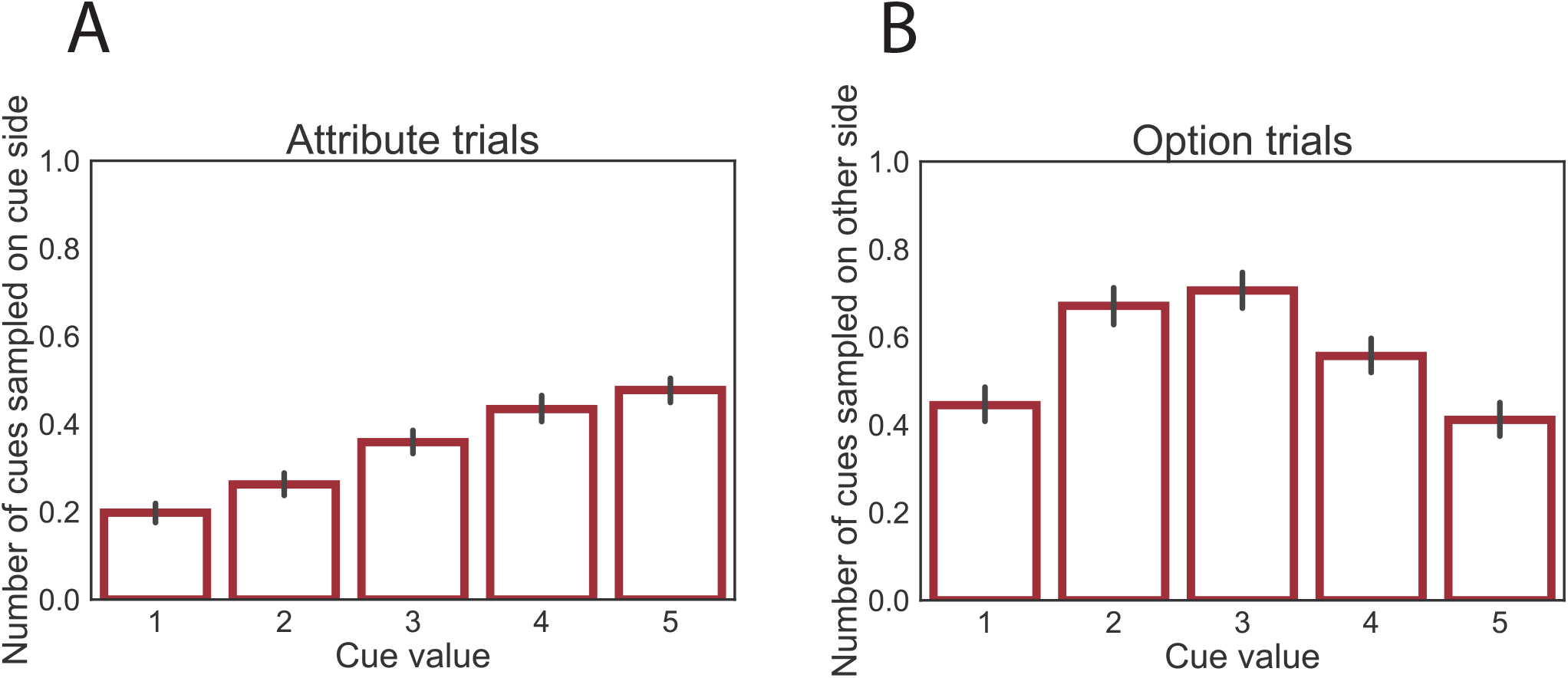
Subsequent information sampling depends on the values of the first two cues. (A) Number of additional cues sampled on the same side as a presented cue as a function of that cue’s value. Participants are more likely to sample another cue from an option if the first cue from that option was of high value. (B) Number of additional cues sampled on opposite side from the presented cue as a function of that cue’s value. Participants are more likely to sample additional cues if the values of the first two cues were of average value.

**Figure S2.**
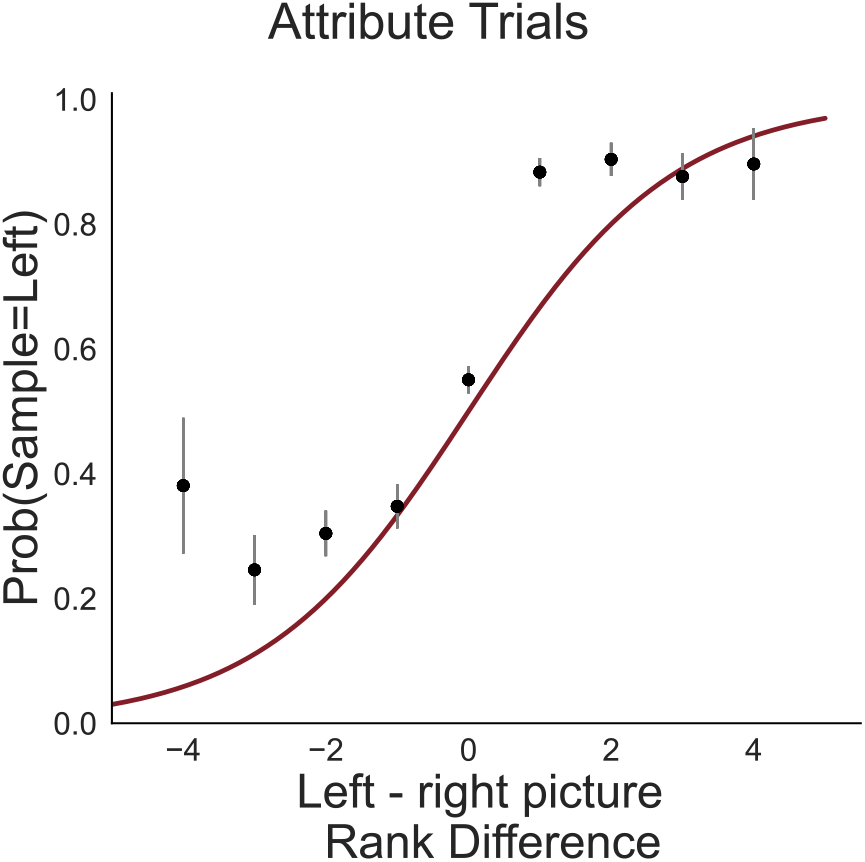
Participants were more likely to sample the remaining hidden cue from the option with the highest revealed cue value in attribute trials. This psychometric curve plots the probability of sampling cues from the left option as a function of the difference in value rank between the two revealed cues and is only plotting trials on which participants sampled additional cues.

**Figure S3.**
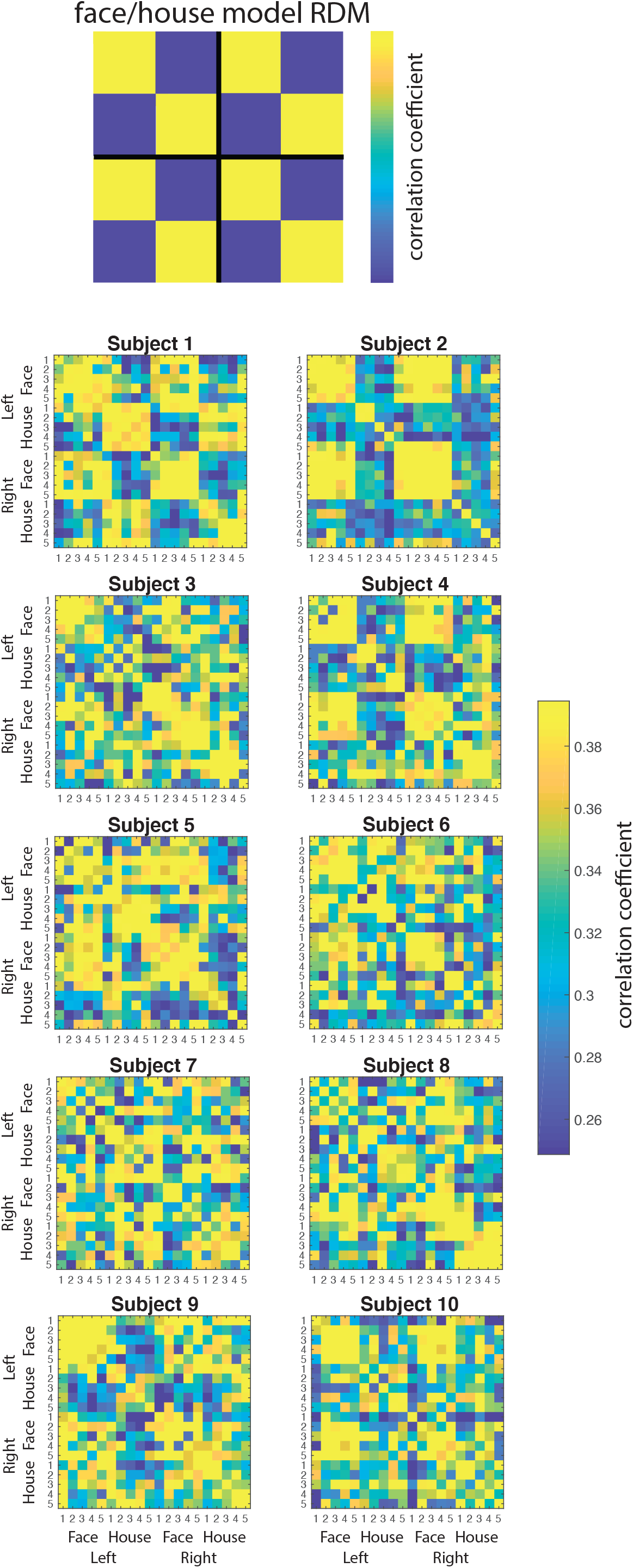
Representational dissimilarity matrices of temporal occipital fusiform gyrus of several example subjects sorted by face/house category and left/right presentation. It can be seen that a clear representation of category is present in almost all participants.

**Figure S4.**
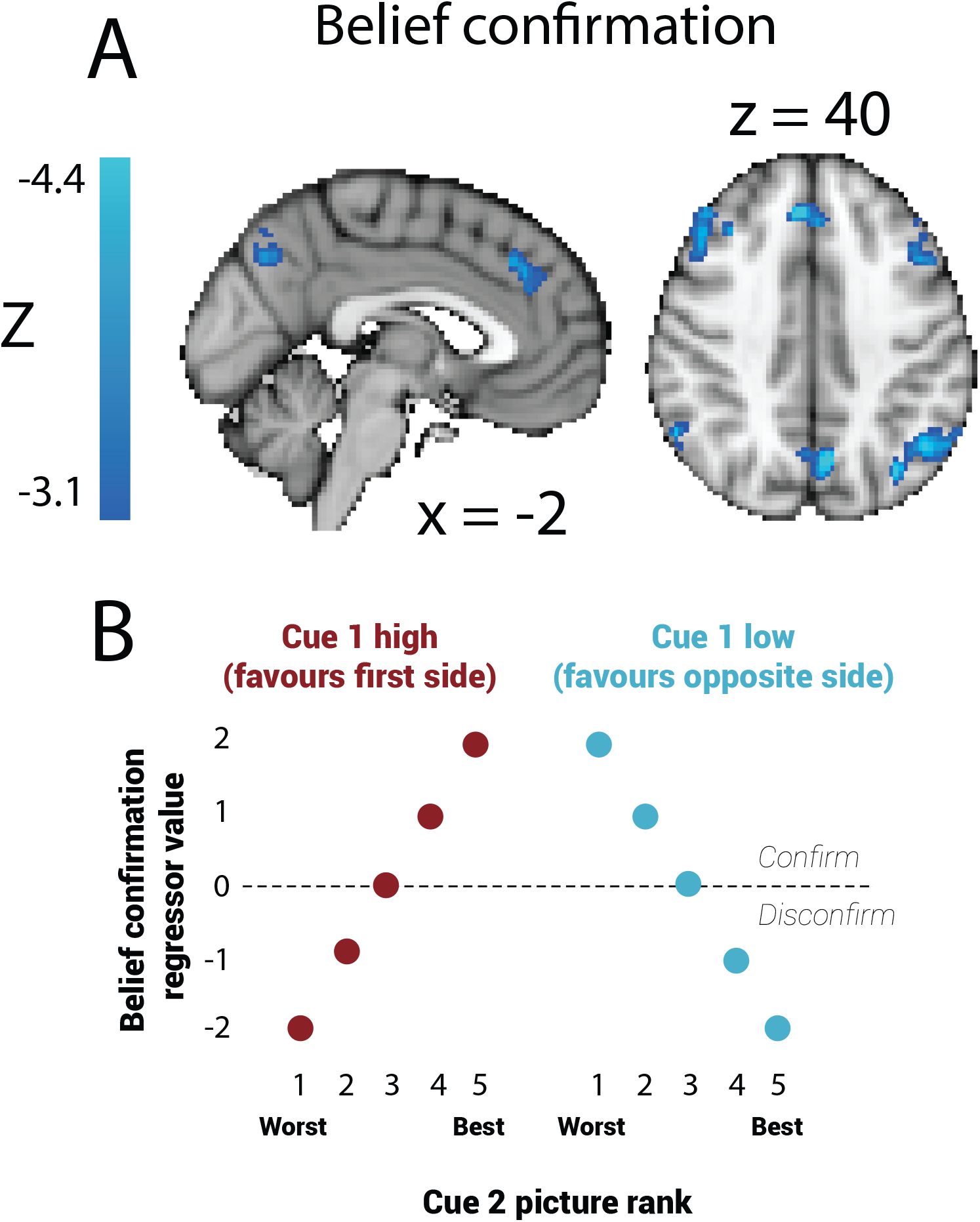
A main effect of belief confirmation was also found in MFC, but this effect disappeared when using the reduced sample of 24 participants. (A) Linear negative effect of belief confirmation in MFC on option trials in 30 subjects. Parametric maps are cluster-corrected and thresholded at z<-3.1. (B) Encoding of belief confirmation as a function of the ranks of the two presented cues.

**Figure S5.**
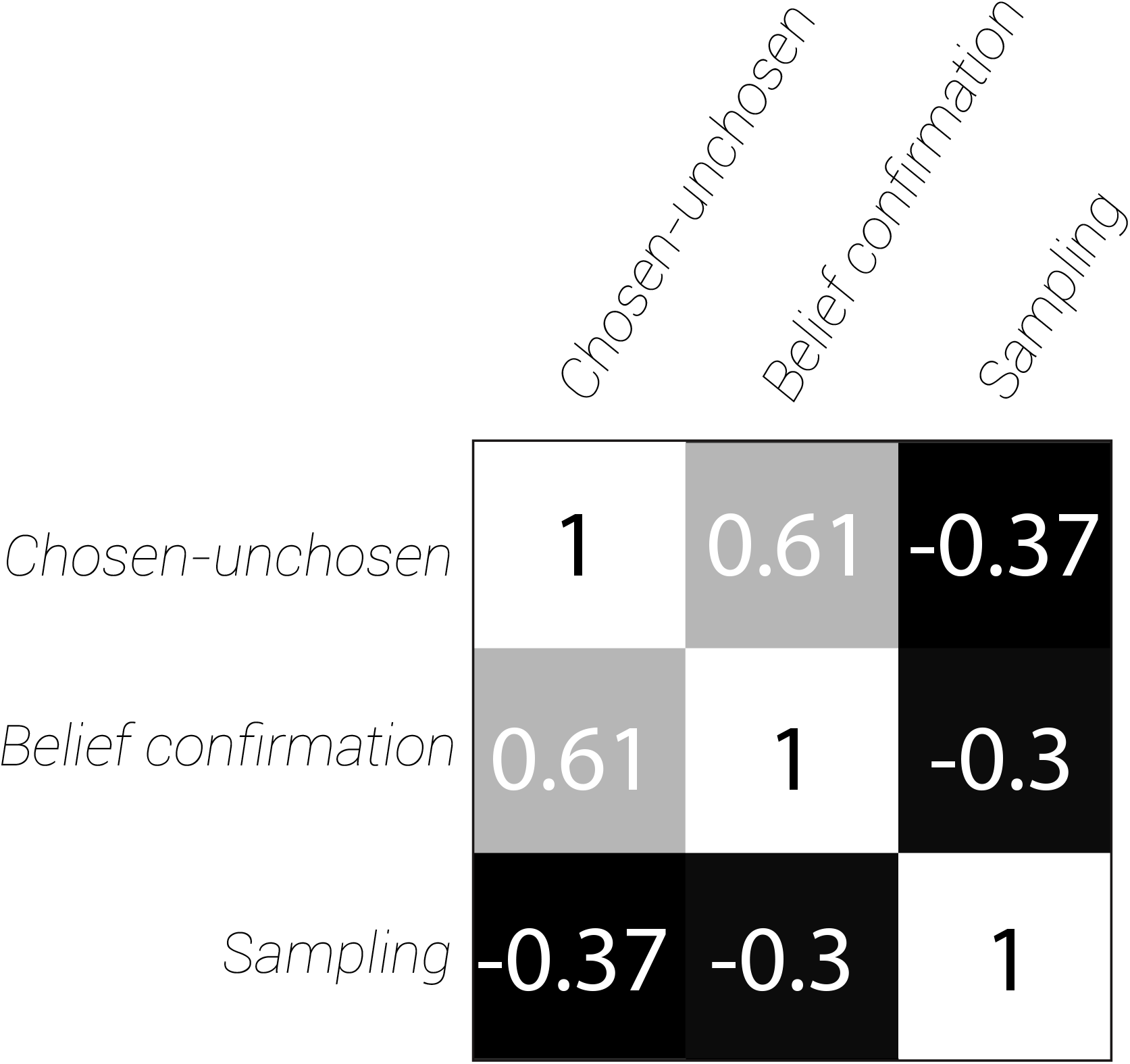
Correlation matrix of value difference, belief confirmation and sampling predictors in mass univariate analysis in 24 subjects.

**Figure S6.**
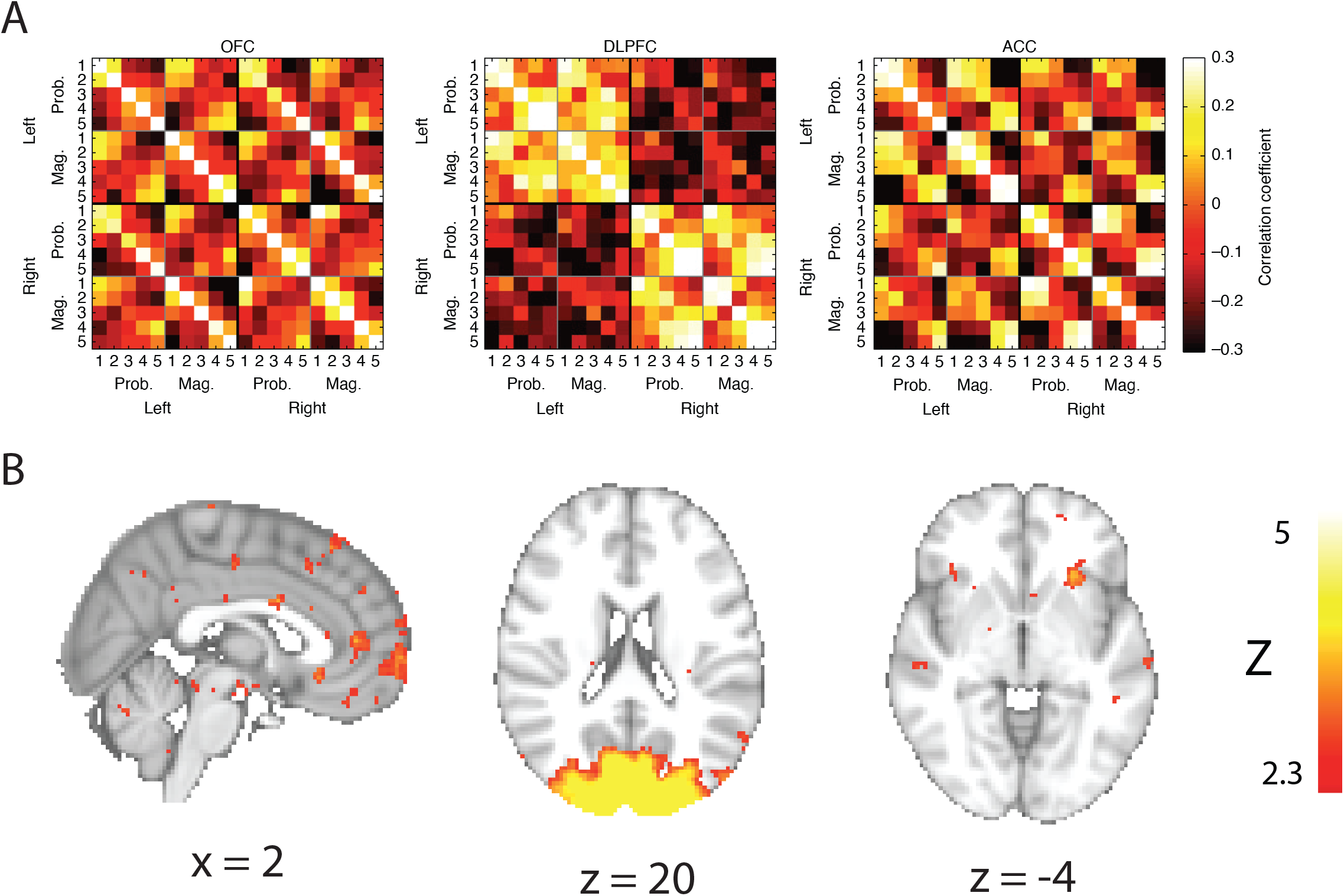
To further investigate RSA as a method to link functional data across species (Kriegeskorte, 2008), we performed a whole-brain searchlight on the human fMRI data using (A) RSA matrices derived directly from the single unit recordings measured in three relevant prefrontal regions in macaques, instead of using model RSA matrices as in Fig. 2 (Hunt et al., 2018). (B) No significant whole-brain cluster-corrected correlations were found within human prefrontal cortex, though subthreshold activations were found in human insula correlated with the macaque ACC matrix, and human ventromedial prefrontal cortex (vmPFC) correlated with the macaque orbitofrontal cortex (OFC) matrix. As expected from the result in Figure 2C/D, human visual cortex was found to be related to the macaque dorsolateral prefrontal cortex (dlPFC) matrix, which strongly reflected the participant’s current locus of spatial attention. All parametric maps are uncorrected and thresholded at z>2.3.

**Figure S7.**
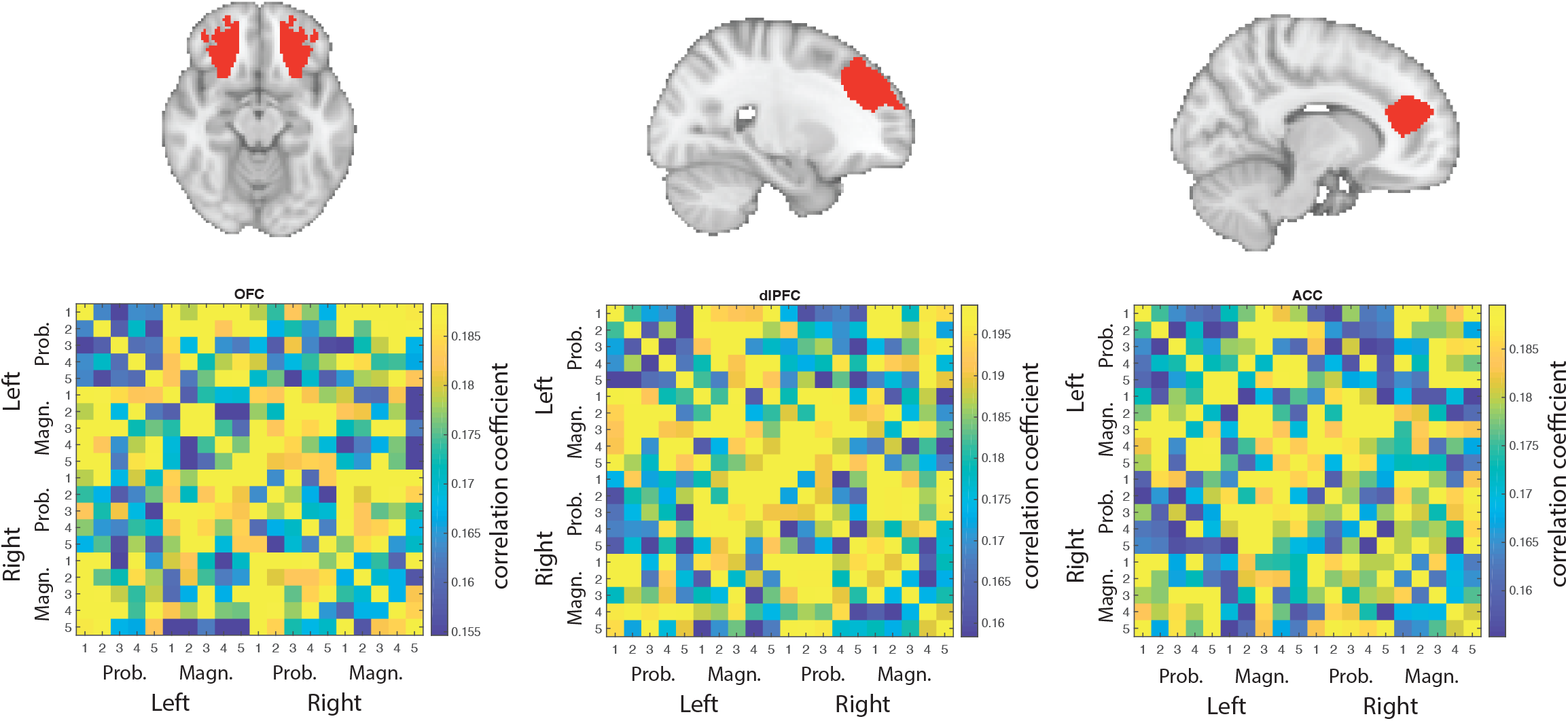
RSA in anatomically defined regions of interest for human orbitofrontal cortex (OFC), dorsolateral prefrontal cortex (dlPFC), and anterior cingulate cortex (ACC) reveals considerably less task-related structure in the representational similarity matrices than those derived from the macaque single-cell data. The ROIs in human PFC are based on connectivity parcellations of human PFC (Sallet et al., 2013; Neubert et al., 2015) and are human homologues of the prefrontal regions studied in Hunt et al. (2018). The matrices are sorted by the 10 possible stimulus identities (five probability cues and five magnitude cues sorted by ranked value) and which side they were presented on. The color of each pixel in the matrix refers to the correlation r between activity in the two conditions across all voxels in the ROI mask.

**Figure S8.**
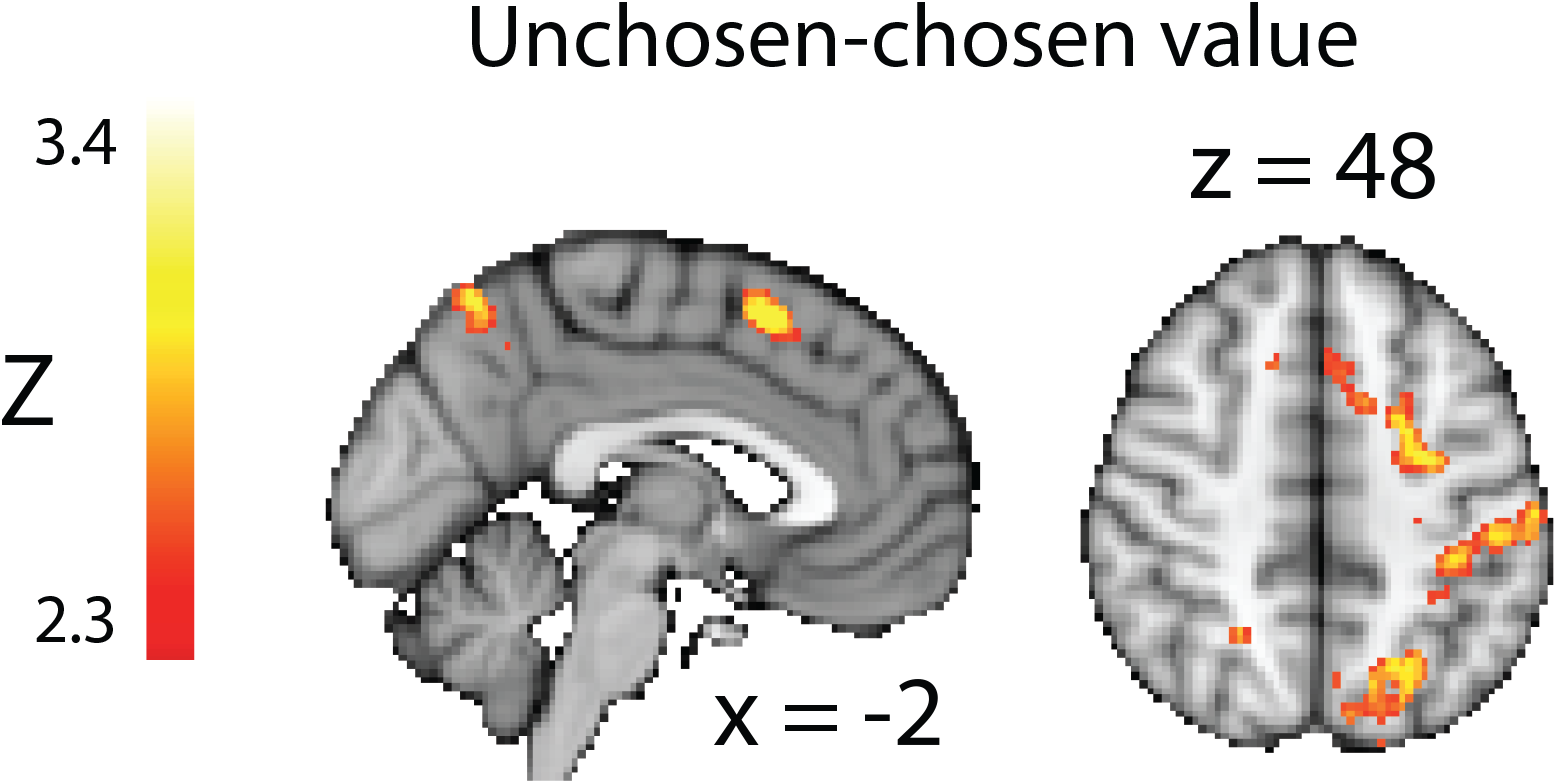
The main effect of inverse value difference at the time of presentation of cue 2 in MFC does not disappear when 5 participants are excluded from sampling. There is a linear positive effect of inverse value difference in MFC on option trials in 24 subjects. Parametric maps are cluster-corrected and thresholded at z>2.3.

**Figure S9.**
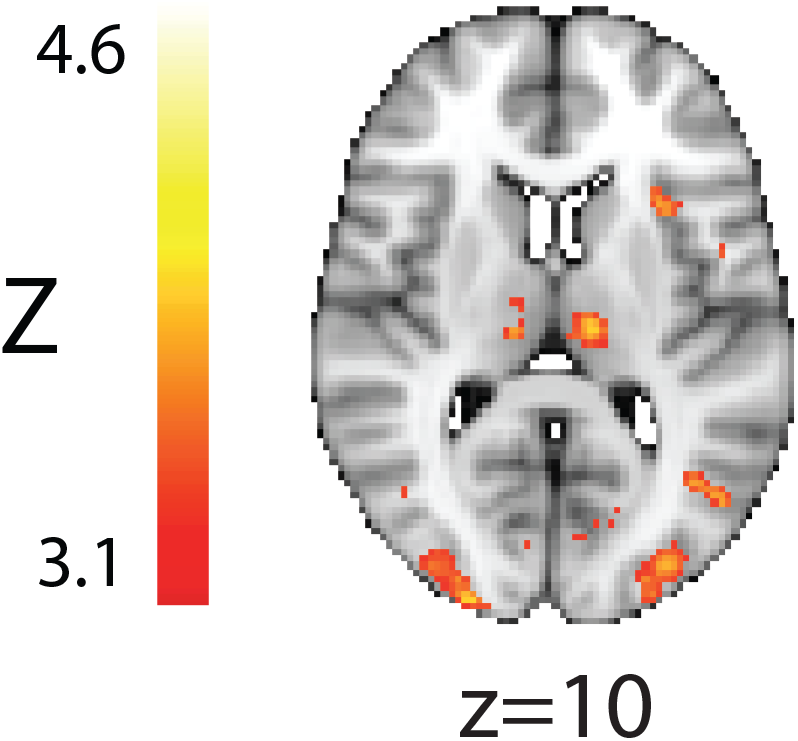
Subthreshold activity found in insula predictive of information sampling in a mass univariate analysis. Parametric map is uncorrected and thresholded at Z>3.1.

